# Comparative genomics of ocular *Pseudomonas aeruginosa* strains from keratitis patients with different clinical outcomes

**DOI:** 10.1101/2020.05.15.097220

**Authors:** Kathirvel Kandasamy, Kannan Thirumalmuthu, Namperumalsamy Venkatesh Prajna, Prajna Lalitha, Vidyarani Mohankumar, Bharanidharan Devarajan

**Affiliations:** Department of Microbiology and Bioinformatics, Aravind Medical Research Foundation, 1 Anna Nagar, Madurai, 625020, Tamil Nadu, India; Department of Cornea, Aravind Eye Hospital, 1 Anna Nagar, Madurai, 625020, Tamil Nadu, India; Department of Clinical Microbiology, Aravind Eye Hospital, 1 Anna Nagar, Madurai, 625020, Tamil Nadu, India; School of Chemical and Biotechnology, SASTRA University, Thanjavur, Tamil Nadu, India

**Keywords:** Bacterial Keratitis, *P. aeruginosa*, whole-genome sequencing, multi-drug resistance, Type III secretion system

## Abstract

Bacterial keratitis caused by *Pseudomonas aeruginosa* is a destructive disease of cornea. Pseudomonas keratitis progresses rapidly and leads to vision loss if untreated. Even with adequate treatment, many patients show poor visual outcomes. The virulence factors or multiple drug-resistant (MDR) mechanisms of the ocular strains responsible for poor clinical outcomes remain largely unknown. Here, we performed whole-genome sequencing of five *P. aeruginosa* strains cultured from corneal scrapings of the healed and corneal buttons of the poor outcome keratitis patients. We investigated the distribution of virulence factors, resistance genes and resistance-associated mutations, the efflux-pump system in all five genomes, and as groups between poor and good clinical outcome as well as MDR vs. non-MDR. We detected several resistance genes and mutations associated with drug resistance in MDR groups; however, a large number of virulence genes were detected in all our genomes. Among the virulence genes, exoU and exoS exotoxin of the Type III secretion system detected in MDR and non-MDR strains, respectively, considered as main virulence contributors of keratitis pathogenesis. Despite this fact, this study did not show an association between MDR with exoU and poor clinical outcomes. However, strain-specific resistance and virulence genes were observed in this study, suggesting their role in the clinical outcome. Mainly, the flagellar genes fliC and fliD, reported to altering the host immune response, might impact the clinical outcome. This comparative study may provide new insights into the genome of ocular strains and requires further functional studies.

## 1. Introduction

*Pseudomonas aeruginosa* is a destructive opportunistic pathogen responsible for bacterial keratitis characterized by ocular pain, infiltration of inflammatory cells, stromal destruction, corneal perforation and leading to vision loss if untreated [1]. Keratitis pathogenesis is a complex process, wherein several virulence factors have been implicated including cell-associated structures such as type IV pili and flagella [2, 3], slime polysaccharide, proteases such as elastase B (LasB), alkaline protease (AprA), protease IV (PrpL) and *P. aeruginosa* small protease (Pasp), and exotoxins [4]. However, several studies revealed that exotoxins exoU and exoS that belong to the type III secretion system (T3SS) is an important contributor to keratitis pathogenesis [5, 6]. *P. aeruginosa* strains expressing the exoS gene are known to invade epithelial cells, whereas strains expressing the exoU gene, a potent phospholipase, are cytotoxic towards host cells [7]. exoU strains has been reported to cause more severe infection [8] than exoS. Moreover, clinical strains from keratitis prefer to carry exoU gene rather than exoS in their accessory genome [5, 9], although the core genome, approximately 90% of the complete genome, contains most of the virulence genes [10]. Besides, the clinical strains carrying multiple antimicrobial drug resistance in the accessory genome are associated with poor clinical outcome in patients with keratitis [11]. Indeed, multiple drug-resistant (MDR) strains carrying exoU protein may lead to poor clinical outcomes [11]. Furthermore, a subpopulation of UK-keratitis associated *P. aeruginosa* strains has been reported to have specific features that cause eye infections [9]. However, the link between specific genomic features and clinical outcome remain unclear.

In this study, we have selected five strains cultured from the keratitis patients based on their drug resistance profiles, the presence of T3SS virulence factors and clinical outcome. First, we report the analysis of annotated genomes of five *P. aeruginosa* ocular strains compared to reference genome PAO1. Next, we study the virulence and drug-resistance mechanisms that impact the clinical outcome.

## 2. Materials and methods

### 2.1. Ocular bacterial strains

Five *P. aeruginosa* strains obtained from keratitis patients attending Aravind Eye Hospital, Madurai, India, were selected retrospectively. The Institutional Review Board approved the study protocol and informed consent was obtained from each patient before sampling. The study was performed according to the tenets of the Declaration of Helsinki. Two strains (BK2 and BK6) cultured from the corneal scrapings of patients who subsequently presented with a healed ulcer after treatment were selected. The other three strains (BK3, BK4 and BK5) cultured from corneal buttons from patients who underwent therapeutic penetrative keratoplastic (TPK) surgery were selected. Bacterial identification was performed based on morphology and growth characteristics in culture, gram’s staining and 16s rDNA sequencing. The strains were stored at −80**°**C at the microbiological culture collection facility.

### 2.2. Antimicrobial Susceptibility Testing

The Minimum Inhibitory Concentration (MIC) of various antibiotics were determined using the broth microdilution method and automated BioMérieux VITEK 2 system following the recommendations of the Clinical and Laboratory Standards Institute (CLSI, 2012). *P. aeruginosa* ATCC^®^ 27853 ™ strain was used for quality control. Briefly, in the broth dilution method, the antibiotic solutions were serially diluted and inoculated with test organisms in Mueller-Hinton broth and incubated for 18-24 hours at 37**°**C. The lowest concentration of an antibiotic that prevented bacterial growth (turbidity) was considered as the MIC. For VITEK based determination, the colonies grown on blood agar for 18-24 hours at 37**°**C were resuspended and adjusted to 0.5 McFarland standard with 0.45% saline. The reagent cards (AST-N281) were inoculated with culture suspensions according to manufacturer’s instructions and data was collected during the entire incubation period.

### 2.3. Type 3 secretion system (T3SS) genotyping

T3SS genotypes were determined by PCR using specific primers (Supplementary Information. SI 3) against exoS, exoT and exoU genes. Briefly, 25 μl PCR reactions were set up with 1x reaction buffer, 200nM primers, 400 μM dNTP and one-unit Taq polymerase. The reaction conditions consisted of an initial denaturation step at 94**°**C for 10min, followed by 35 cycles of 94**°**C for 2min, 60**°**C for 30s and 72**°**C for 1 min, with a final extension at 72**°**C for 5min. The reference strains PAO1 and PA14 served as positive controls.

### 2.4. DNA isolation and genome sequencing

Genomic DNA was isolated from strains using the QIAamp DNA mini kit from Qiagen (Hilden, Germany) according to their manufactures protocol. NanoDrop spectrophotometer (Thermo Scientific, Wilmington, USA) was used for purity and concentration of DNA. Whole-genome sequencing (WGS) was performed at Genotypic, India. Briefly, the genomic DNA was fragmented to obtain 200 to 600 bp size by Covaris S220. Paired-end libraries (∼140-480bp insert size) were prepared using NEXTFlex DNA sequencing kit and quantified with Bioanalyzer (Agilent). WGS was performed using Illumina NextSeq 500 platform and the raw data of each strain were obtained through de-multiplexing.

### 2.5. Genome assembly and annotation

The raw data was quality checked and preprocessed to remove the adapter sequences, low coverage reads and PCR duplicates. The reads were further assembled into contigs with CLC genomic workbench (v9.0.1). Contigs ordering and visualizations were performed using Mauve program v2.3.1 [12] using PAO1 genome as a reference. The assembly was further improved using Genome Finishing Module in the CLC genomics workbench. Rapid Annotations Subsystems Technology (RASTtk) [13] server was used for annotation.

### 2.6. Comparative genomics

Mauve and Roary v.3.12.0 [14] was used to perform comparative genomics and pangenome analysis. MLST prediction server [15] was used to identify sequence types of each strain. ResFinder [16] was used for the detection of antibiotic resistance genes and 90 % threshold and 80 % of minimum length were set. Virulence Factor of Pathogenic Bacteria database (VFDB) [17] with E value cut-off of 1e-5 was used for virulence gene identification and PHAST server for prophage sequence identification [18]. Clustered Regularly Interspaced Short Palindromic Repeats (CRISPR) and spacer sequences were identified by searching the genome sequence of five strains against CRISPRCasFinder [19].

### 2.7. Validation by Sanger Sequencing

Bi-directional sequencing method was performed to validate the mutations in genes that are associated with antibiotic resistance. The primers sequences, annealing temperatures, and amplicon sizes are mentioned in Supplementary Table 2. A thermal gradient PCR was performed with 120nM primers, 400 μM dNTP and one unit Taq polymerase in a 25 μl reaction. 3130 Genetic Analyzer (Applied Biosystems) was used for sequencing.

### 2.8. Motility Assay

*Swimming:* Swim plates (1% tryptone, 0.5% NaCl, 0.3% agar) were inoculated with a 0.2 μl of overnight cultures and incubated for 24 h at 30°C. The turbid zone formed by the bacterial cells migrating away from the point of inoculation was assessed. *Swarming:* 0.2 μl of overnight cultures were spotted on the surface of the swarm plates (0.5% Luria Bertani (LB) agar and 0.5% glucose) and incubated for 24 h at 30°C. *Twitching:* 1 μl of overnight cultures were stabbed into the 1.5% LB agar plates until the culture reaches the surface of the plate. After 24 hrs of incubation at 37°C, a hazy zone of growth at the interface between the agar and the polystyrene surface of the plate was assessed by the crystal violet staining (1% [wt/vol] solution) after the removal of the agar. PAO1 and PA14 were used as control.

### 2.9. Data Access

The whole-genome sequencing data is available at NCBI-SRA with accession number SRR2062214, SRR2063971, SRR2063976, SRR2063980 and SRR2063985.

## 3. Results and Discussion

### 3.1. Antibiotic susceptibility profile and Genome characteristics

BK3 and BK6 strains were identified as MDR strains (showed resistance to more than three class of antibiotics), whereas BK2, BK4 and BK5 were identified as non-MDR as shown in Table 1. All the strains were sensitive to Colistin. WGS yielded 2.3 - 3.2 GB of data from all five strains. An average of 65.5% of GC content was observed (Table 2). MDR strains were found to have a bigger genome size of 7.1 MB than non-MDR strains, consisting of more than 100 scaffolds. Also, more genes were detected in MDR strains. However, the number of tRNA genes was similar among them. Further, we constructed a phylogenetic tree using core SNPs detected by Parsnp tool [20] in *P. aeruginosa* genomes from keratitis patients. We included our genomes, previously reported genomes from India, Australia [11, 21], and England [9], and PAO1 and PA14 (Fig. 1). We observed that our MDR strains closely related to Indian (PA31, PA32, PA33, PA35 and PA37) and England strain (39016), while our non-MDR strains were closely related to Australian strains (PA17, PA40, PA149, PA157 and PA175). Nevertheless, we observed that the strains were distantly related based on their clinical outcome.

**Fig. 1.**
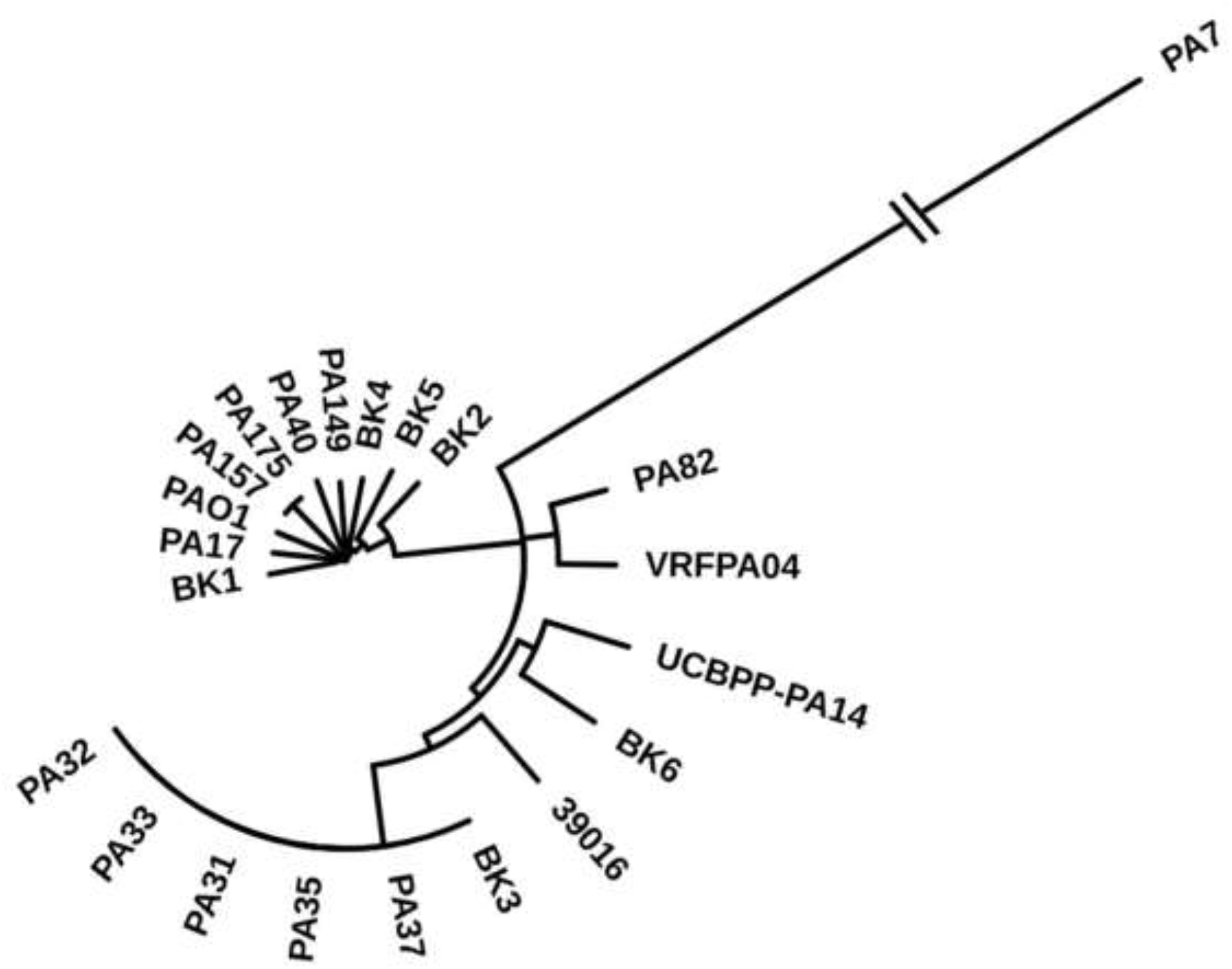
Phylogenetic tree of *P. aeruginosa* strains. Maximum likelihood tree was created based on core SNPs mapped with PAO1 as reference genome.

**Table 1.**
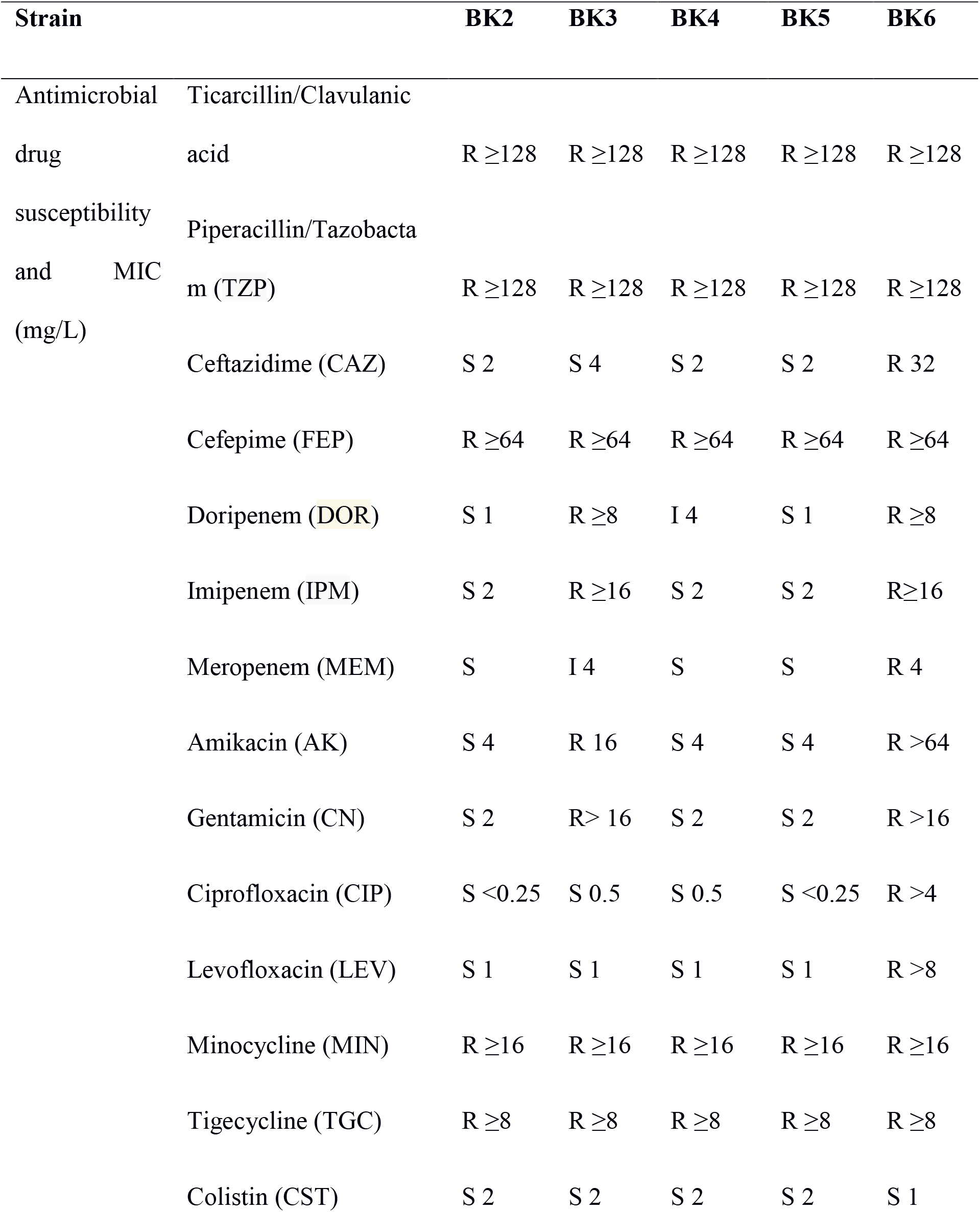
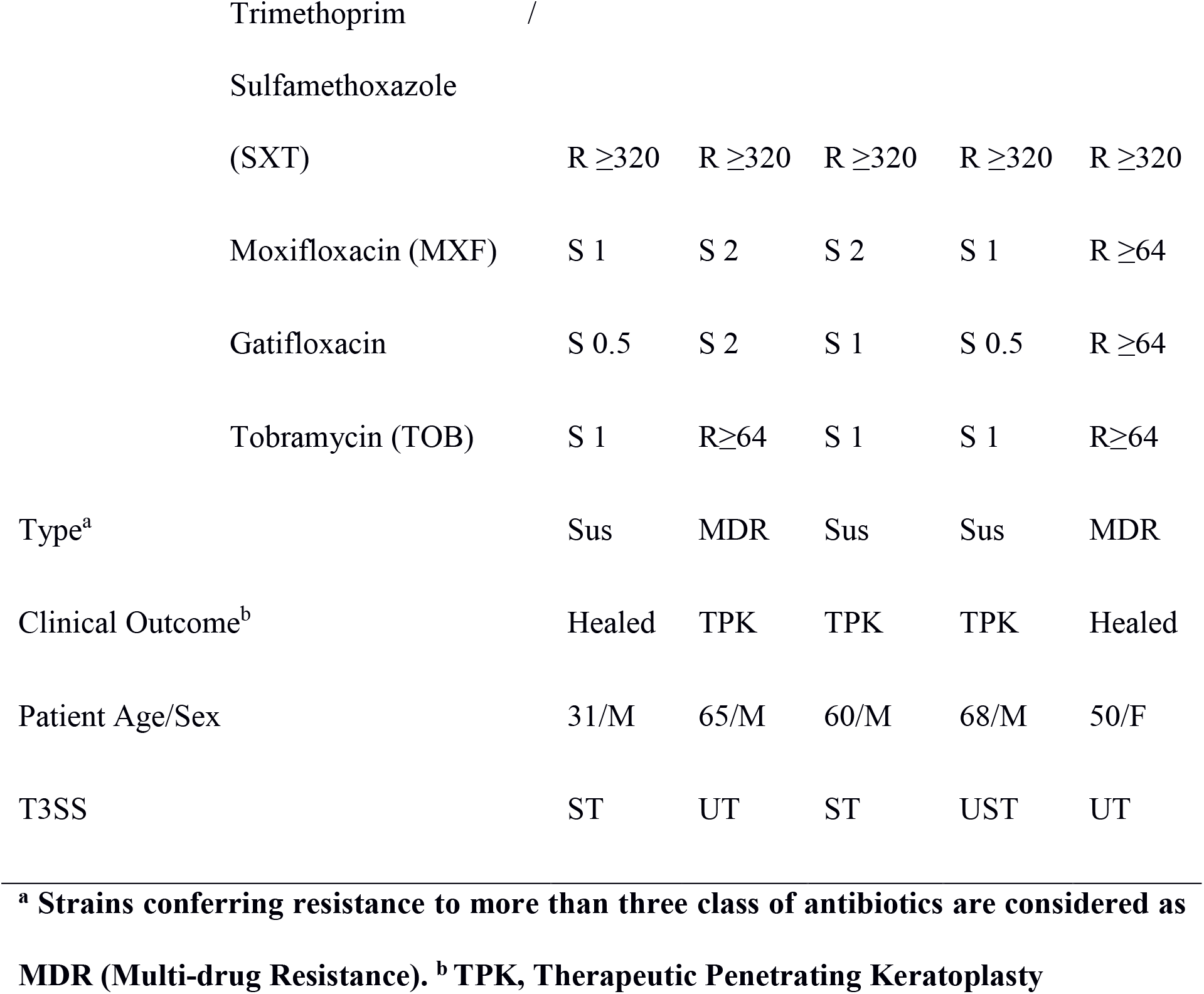
Minimum inhibitory concentration and clinical data of *P. aeruginosa* ocular strains. R, resistant; I, intermediate; S, susceptible.

| Strain |  | BK2 | BK3 | BK4 | BK5 | BK6 |
| --- | --- | --- | --- | --- | --- | --- |
| Antimicrobial drug susceptibility and MIC (mg/L) | Ticarcillin/Clavulanic acid | R ≥128 | R ≥128 | R ≥128 | R ≥128 | R ≥128 |
| | Piperacillin/Tazobactam (TZP) | R ≥128 | R ≥128 | R ≥128 | R ≥128 | R ≥128 |
|  | Ceftazidime (CAZ) | S 2 | S 4 | S 2 | S 2 | R 32 |
| | Cefepime (FEP) | R ≥64 | R ≥64 | R ≥64 | R ≥64 | R ≥64 |
| | Doripenem (DOR) | S 1 | R ≥8 | I 4 | S 1 | R ≥8 |
| | Imipenem (IPM) | S 2 | R ≥16 | S 2 | S 2 | R ≥16 |
|  | Meropenem (MEM) | S | I 4 | S | S | R 4 |
|  | Amikacin (AK) | S 4 | R 16 | S 4 | S 4 | R >64 |
|  | Gentamicin (CN) | S 2 | R > 16 | S 2 | S 2 | R >16 |
|  | Ciprofloxacin (CIP) | S <0.25 | S 0.5 | S 0.5 | S <0.25 | R >4 |
|  | Levofloxacin (LEV) | S 1 | S 1 | S 1 | S 1 | R >8 |
| | Minocycline (MIN) | R ≥16 | R ≥16 | R ≥16 | R ≥16 | R ≥16 |
| | Tigecycline (TGC) | R ≥8 | R ≥8 | R ≥8 | R ≥8 | R ≥8 |
|  | Colistin (CST) | S 2 | S 2 | S 2 | S 2 | S 1 |

|  |  |  |  |  |  |
| --- | --- | --- | --- | --- | --- |
| Trimethoprim / Sulfamethoxazole (SXT) | R ≥320 | R ≥320 | R ≥320 | R ≥320 | R ≥320 |
| Moxifloxacin (MXF) | S 1 | S 2 | S 2 | S 1 | R ≥64 |
| Gatifloxacin | S 0.5 | S 2 | S 1 | S 0.5 | R ≥64 |
| Tobramycin (TOB) | S 1 | R ≥64 | S 1 | S 1 | R ≥64 |
| Type<sup>a</sup> | Sus | MDR | Sus | Sus | MDR |
| Clinical Outcome<sup>b</sup> | Healed | TPK | TPK | TPK | Healed |
| Patient Age/Sex | 31/M | 65/M | 60/M | 68/M | 50/F |
| T3SS | ST | UT | ST | UST | UT |

<sup>a</sup> Strains conferring resistance to more than three class of antibiotics are considered as MDR (Multi-drug Resistance). <sup>b</sup> TPK, Therapeutic Penetrating Keratoplasty

**Table 2.** Genomic features and statistics of ocular *P. aeruginosa* strains.

| **Strain** | **BK2** | **BK3** | **BK4** | **BK5** | **BK6** |
| --- | --- | --- | --- | --- | --- |
| Genome size (Mb) | 6.3 | 7.1 | 6.5 | 6.3 | 7.1 |
| G+C | 66 % | 65 % | 64 % | 66 % | 66 % |
| Scaffolds | 45 | 113 | 69 | 68 | 129 |
| Maximum length (bp) | 703333 | 594831 | 700326 | 877303 | 602407 |
| Minimum length (bp) | 506 | 504 | 509 | 509 | 510 |
| N50 | 370564 | 191913 | 347365 | 350169 | 232253 |
| CDs | 6037 | 7009 | 6141 | 6055 | 6722 |
| Hypothetical proteins | 1042 | 1460 | 1079 | 1078 | 1327 |
| tRNA | 61 | 65 | 60 | 61 | 60 |

### 3.2. Pattern of gene distribution

Nine thousand two orthologs were detected in our five genomes and PAO1 compared to 9344 orthologs of reported *P. aeruginosa* genomes [22]. Among the detected orthologs, we identified 5139 genes as core genome. While in the accessory genome of each strain, we identified 165, 551, 161, 162, 429 and 115 in BK2, BK3, BK4, BK5, BK6 and PAO1 respectively as unique (strain-specific) genes. Further, we checked Clusters of Orthologous Groups (COG) of core and unique genes (Supplementary Fig. S1). The core genome was enriched to class E (Amino acid transport and metabolism) and class T (Signal transduction mechanisms). The unique genes were enriched to class L (Replication, recombination and repair), class M (Cell wall/membrane/envelope biogenesis), class I (Lipid transport and metabolism) and class S (Function unknown), suggesting that they may help for bacterial survival.

### 3.3. Antibiotic Resistance

#### 3.3.1. Resistance genes

We detected twenty-one antibiotic resistance genes in all five genomes. Seven genes were present in all five, as shown in Table 3. Among them, the β-lactamase genes blaPAO and blaOXA-50, are known to cause resistance to the carbapenem class of antibodies [23, 24]. The presence of aph(3’)-IIb gene may provide resistance to many aminoglycoside class of antibiotics, pmrA-pmrB gene may provide resistance to polymyxin B and fosA gene to Fosfomycin. The catB7 gene of CATB family was reported to cause chloramphenicol resistance [25]. As expected, we observed more genes (17) in MDR than non-MDR strains (11). Besides, both groups carried several strain-specific genes; 16S rRNA methylase genes (rmtA and rmtB), aminoglycoside acetyltransferase gene (aac(3)-Id), blaVIM, rosB and drfB5 in BK6, streptomycin-inactivating enzymes (strA and strB) and sul1 in BK3, and PDC-5 in BK5. The tetG gene, responsible for tetracycline resistance was detected in both MDR strain. The strain-specific genes detected in MDR strains were not reported earlier in other ocular *P. aeruginosa* genomes [9, 21]. Indeed, our previous study has reported that aminoglycoside resistance among the ocular *P. aeruginosa* strains was mediated mainly through the enzymatic modification of drugs [26].

**Table 3.**
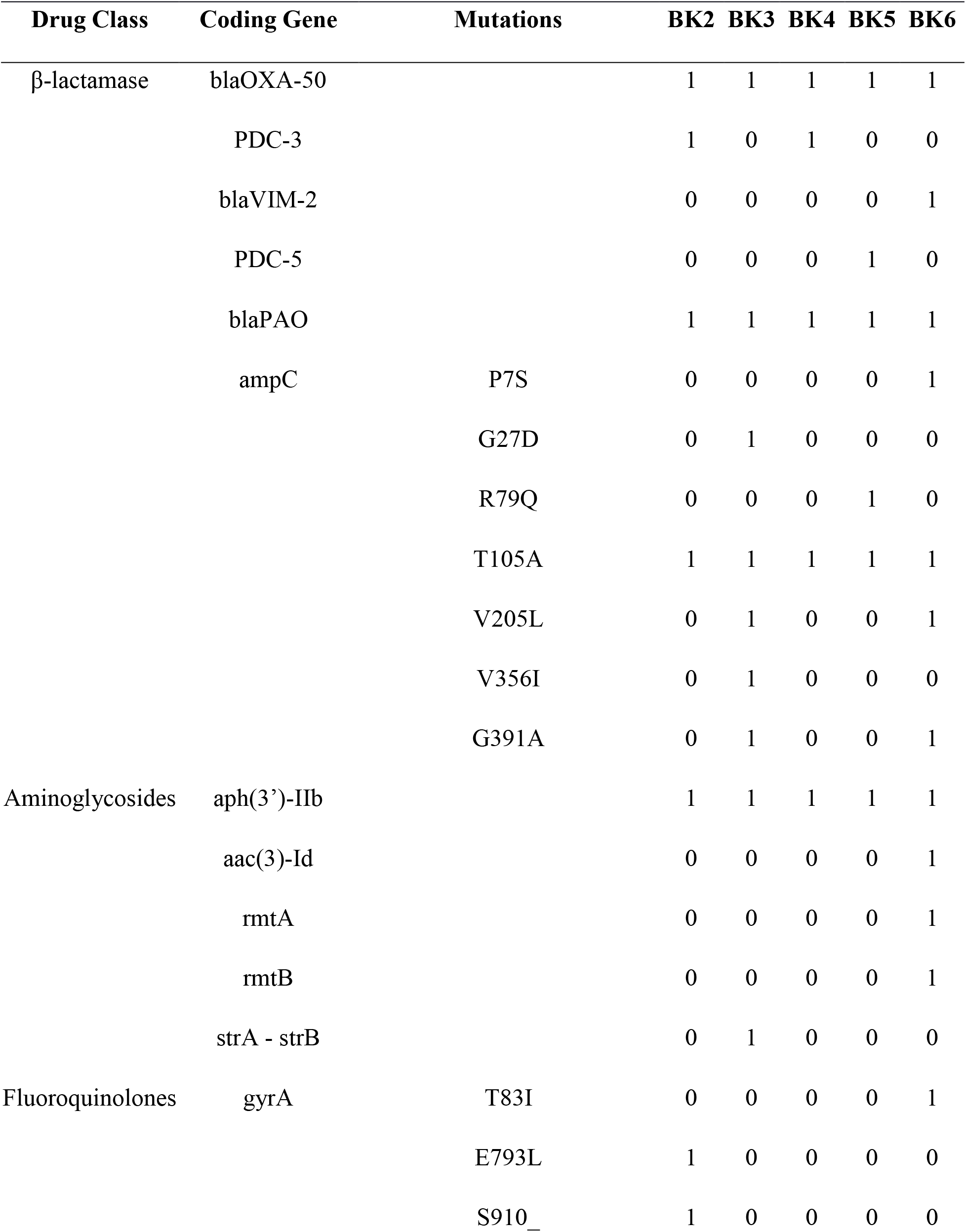
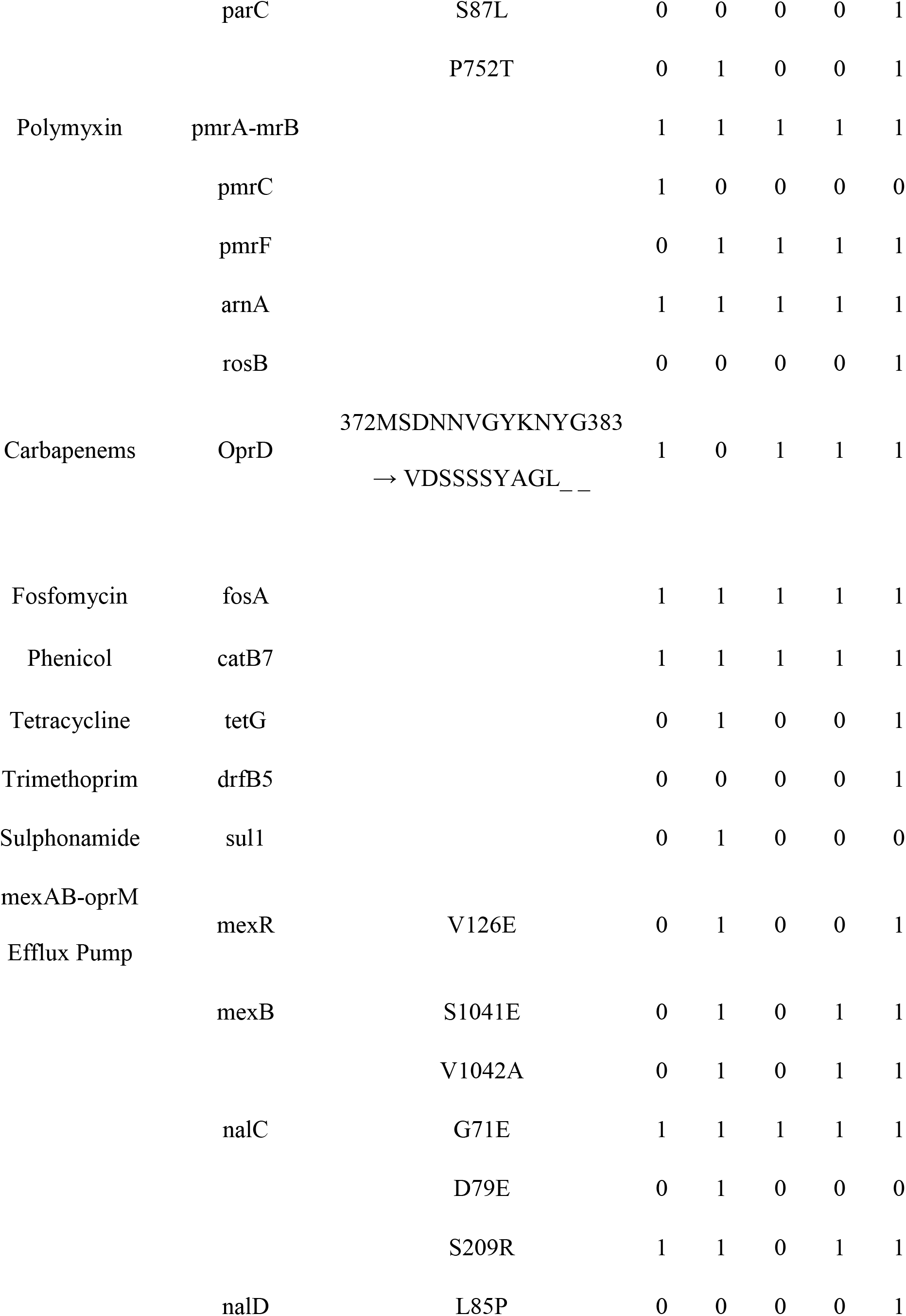
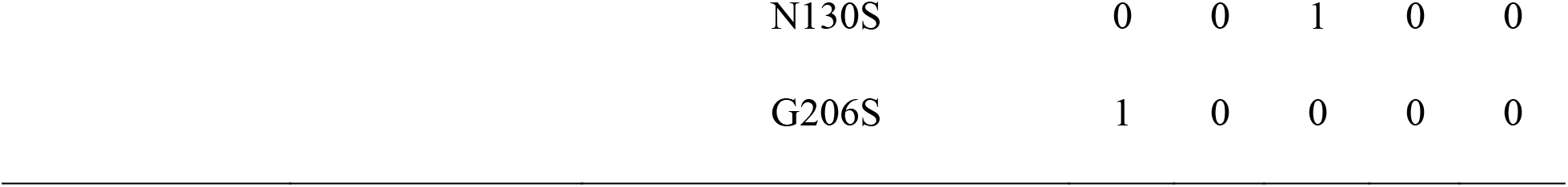
Antimicrobial resistance-associated genes and resistance-associated mutations detected in all five ocular strains of *P. aeruginosa* compared to PAO1. 1 = Present, 0 = Absent.

| Drug Class | Coding Gene | Mutations | BK2 | BK3 | BK4 | BK5 | BK6 |
| --- | --- | --- | --- | --- | --- | --- | --- |
| β-lactamase | blaOXA-50 | | 1 | 1 | 1 | 1 | 1 |
|  | PDC-3 |  | 1 | 0 | 1 | 0 | 0 |
|  | blaVIM-2 |  | 0 | 0 | 0 | 0 | 1 |
|  | PDC-5 |  | 0 | 0 | 0 | 1 | 0 |
|  | blaPAO |  | 1 | 1 | 1 | 1 | 1 |
|  | ampC | P7S | 0 | 0 | 0 | 0 | 1 |
|  |  | G27D | 0 | 1 | 0 | 0 | 0 |
|  |  | R79Q | 0 | 0 | 0 | 1 | 0 |
|  |  | T105A | 1 | 1 | 1 | 1 | 1 |
|  |  | V205L | 0 | 1 | 0 | 0 | 1 |
|  |  | V356I | 0 | 1 | 0 | 0 | 0 |
|  |  | G391A | 0 | 1 | 0 | 0 | 1 |
| Aminoglycosides | aph(3')-IIb |  | 1 | 1 | 1 | 1 | 1 |
|  | aac(3)-Id |  | 0 | 0 | 0 | 0 | 1 |
|  | rmtA |  | 0 | 0 | 0 | 0 | 1 |
|  | rmtB |  | 0 | 0 | 0 | 0 | 1 |
|  | strA - strB |  | 0 | 1 | 0 | 0 | 0 |
| Fluoroquinolones | gyrA | T83I | 0 | 0 | 0 | 0 | 1 |
|  |  | E793L | 1 | 0 | 0 | 0 | 0 |
|  |  | S910_ | 1 | 0 | 0 | 0 | 0 |
|  | parC | S87L | 0 | 0 | 0 | 0 | 1 |
|  |  | P752T | 0 | 1 | 0 | 0 | 1 |
| Polymyxin | pmrA-mrB |  | 1 | 1 | 1 | 1 | 1 |
|  | pmrC |  | 1 | 0 | 0 | 0 | 0 |
|  | pmrF |  | 0 | 1 | 1 | 1 | 1 |
|  | arnA |  | 1 | 1 | 1 | 1 | 1 |
|  | rosB |  | 0 | 0 | 0 | 0 | 1 |
| Carbapenems | OprD | 372MSDNNVG YK NYG383<br>→ VDSSSSYAGL__ | 1 | 0 | 1 | 1 | 1 |
| Fosfomycin | fosA |  | 1 | 1 | 1 | 1 | 1 |
| Phenicol | catB7 |  | 1 | 1 | 1 | 1 | 1 |
| Tetracycline | tetG |  | 0 | 1 | 0 | 0 | 1 |
| Trimethoprim | drfB5 |  | 0 | 0 | 0 | 0 | 1 |
| Sulphonamide | sul1 |  | 0 | 1 | 0 | 0 | 0 |
| mexAB-oprM<br>Efflux Pump | mexR | V126E | 0 | 1 | 0 | 0 | 1 |
|  | mexB | S1041E | 0 | 1 | 0 | 1 | 1 |
|  |  | V1042A | 0 | 1 | 0 | 1 | 1 |
|  | nalC | G71E | 1 | 1 | 1 | 1 | 1 |
|  |  | D79E | 0 | 1 | 0 | 0 | 0 |
|  |  | S209R | 1 | 1 | 0 | 1 | 1 |
|  | nalD | L85P | 0 | 0 | 0 | 0 | 1 |

|  |  |  |  |  |  |
| --- | --- | --- | --- | --- | --- |
| N130S | 0 | 0 | 1 | 0 | 0 |
| G206S | 1 | 0 | 0 | 0 | 0 |

#### 3.3.2. Gene mutations associated with drug resistance

To identify drug resistance-associated point mutations leading to amino acid change, we detected mutations in drug target genes or mutations that overexpress resistance protein using PAO1 as reference (Table 3). We detected several mutations in ampC gene. Among them, G27D, R79Q, T105A, V205L and G391A were reported to overexpress intrinsic ampC protein [27, 28], and thus they provide penicillin and cephalosporin resistance to ocular strains. Of these, R79Q and G27D were detected only in BK5 and BK3, respectively. Also, we identified mutations in fluoroquinolones target genes gyrA (DNA gyrase) and parC (Topoisomerase IV). A gyrA mutation (T83I) detected in BK6 was previously reported to cause resistance to fluoroquinolones [29, 30]. In addition, three strain-specific gyrA mutations detected in BK2 and one parC mutation (S87L) in BK6 were only observed in this study. Studies report that mutations in gyrA along with mutations in parC might provide the higher-level resistance to fluoroquinolones [29, 31]. Indeed, we previously observed that T83I in gyrA and S87L in parC were predominant amino acid substitutions [26]. Interestingly, we detected oprD gene mutation that leads to twelve amino acid alterations in all strains except BK3. The oprD gene is an outer membrane porin that facilitates the uptake of carbapenem drugs in *P. aeruginosa* [32], and thus the oprD mutations confer the resistance to carbapenems, especially to imipenem.

Genes of multiple efflux pump systems, MexAB-OprM, MexCD-OprJ and MexEF-OprN, which belongs to RND (resistance-nodulation-division) efflux pump family, was identified in our strains. However, we detected non-synonymous point mutations only in transcriptional regulator genes mexR, nalC and nalD that govern the expression of MexAB-OprM pump (Table 3). A mexR gene mutation that leads to V126E detected in MDR strains BK3 and BK6, reported to overexpress the MexAB-oprM pump and thus increase the resistance to fluoroquinolones [29]. Also, nalC mutations, G71E and S209R were detected in all strains, while D79E detected only in BK3. Interestingly, three novel mutations L85P, N130S and G206S of nalD, a secondary repressor of the MexAB-OprM system, were detected in BK6, BK3 and BK2, respectively. A study showed that mutations in the nalD gene might overexpress mexAB-OprM efflux [33].

#### 3.3.3. Virulence factors

We detected several virulence factors including T3SS, Type IV Pili and Fimbrial biogenesis, Pyoverdin biosynthesis, Phenazine biosynthesis, Alginate regulation, Flagellar biosynthesis, Paerucumarin biosynthesis, lasA protease precursor, Twitching motility and Type VI Secretion System in all five genomes (Fig. 2). Among the T3SS toxins, gene encoding exoU was identified in BK3, BK5 and BK6, while exoS, exoT and exoY genes were present in all the strains. However, the exoS was not predicted to be functional in BK3 and BK6 due to a very low identity with functional exoS (Fig.2). Interestingly, a single nucleotide insertion in exoU gene was detected in BK5, which leads to a non-sense mutation (p.Q511X) and possibly truncating 163 amino acids at the C-terminal domain. Therefore, we report here that all MDR strains carry exoU and non-MDR strains carry exoS for their virulence.

**Fig. 2.**
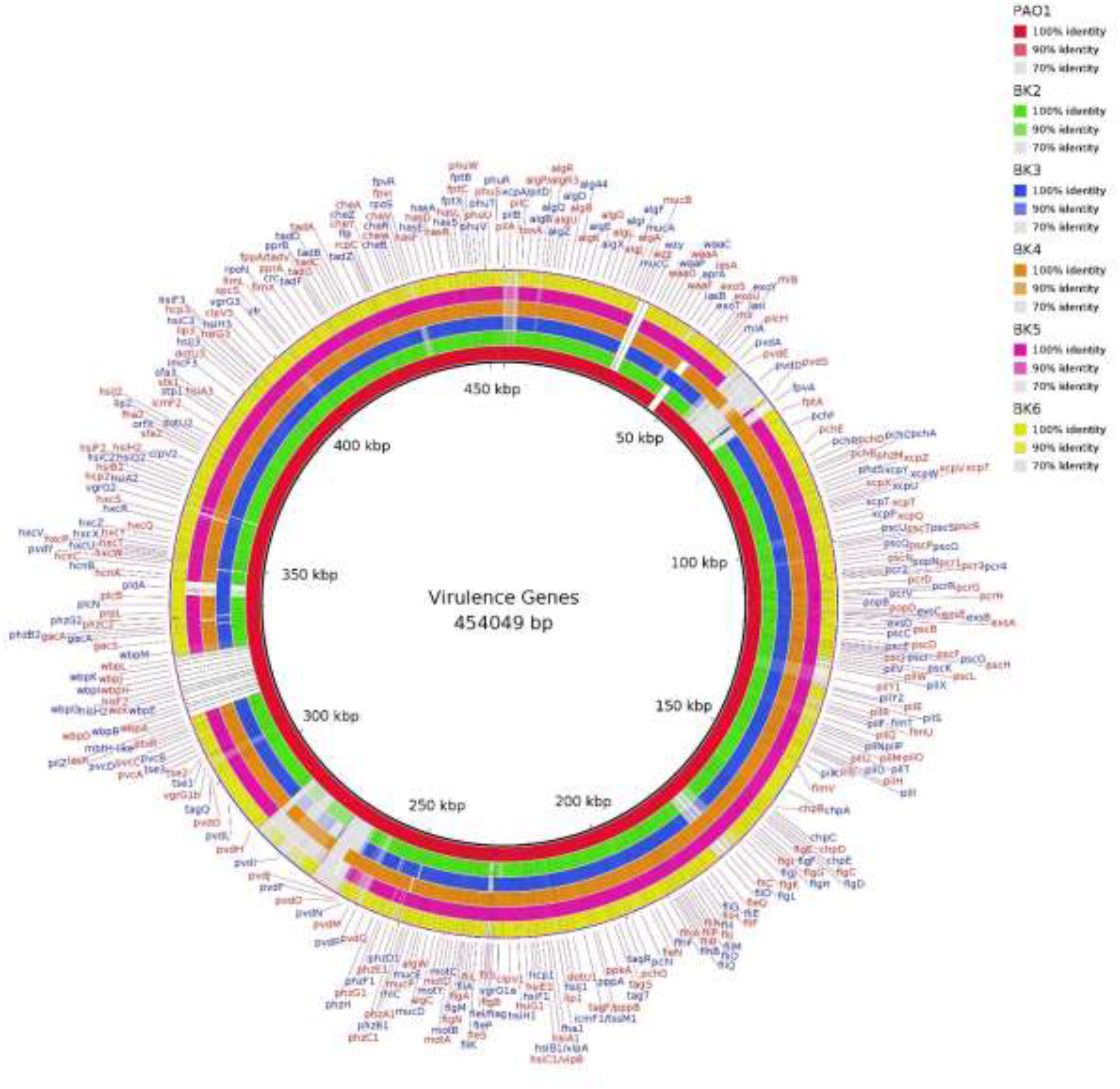
The virulence genes detected in ocular *P. aeruginosa* strains and PAO1. The color coding indicates the sequence identity in percentage against VFDB database.

Among the strains from the poor clinical outcome group, the flagellar genes (fliC, fliD, fliS, flgL, fleP, flgK and fleI/flag) and pyoverdine synthetase gene (pvdY) were detected with more than 90% identity with PAO1 in BK4 and BK5. Of these flagellar genes, fliC and fliD has been reported to induce inflammasome and impairs bacterial clearance [34]. Furthermore, flagellar genes are essential for swimming, twitching and swarming motility for bacteria [35, 36]. We performed motility assay in our ocular strains, PAO1 and PA14. Both BK4 and BK5 strains showed all three motility (Fig. 3). Besides, BK2 also showed all three motilities, suggesting that it might have other motility genes. Indeed, the pilB and pilC genes detected in BK2 with more than 90% sequence identity compared to other strains, were essential for type IV pilus (T4P) expression and twitching motility [37, 38].

**Fig. 3.**
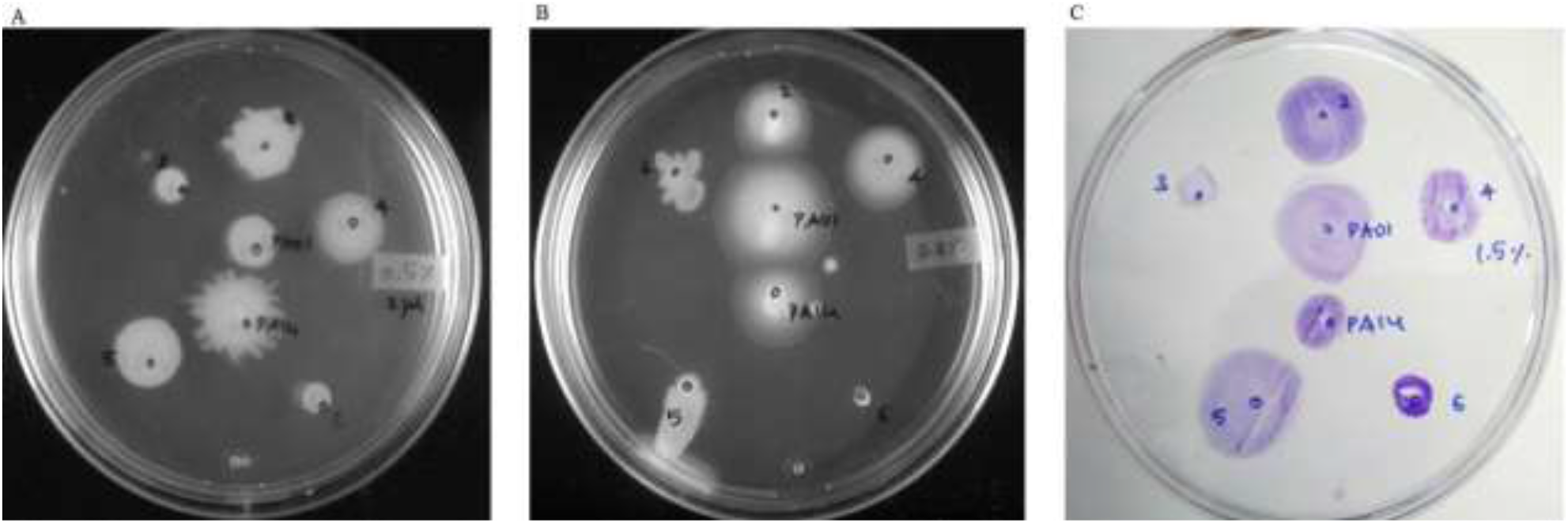
Motility assays A. swarming B. swimming and C. twitching for *P. aeruginosa* ocular strains and reference strains PAO1 and PA14. The numbers in the culture plates 2-6 represents the ocular strains BK2-BK6 respectively.

#### 3.3.4. Prophages

The f1-like bacteriophages, a family of Inoviridae that contribute to bacterial short term evolution and virulence in *P. aeruginosa* [39] were found in BK2, BK4 and BK6 strains, (Supplementary Table 1). A cytotoxin converting P2-like phage of *P. aeruginosa,* phiCTX was identified in BK4, BK5 and BK6. A lytic E. Coli phage vB_EcoM_ECO1230-10, which belongs to Myoviridae family and Mu-like transposable bacteriophage B3 were identified in BK2. F10, a *Pseudomonas* temperate phage belongs to siphovirus family, was found in BK3. Also, we detected strain-specific phages in genomes of poor clinical outcome group that include Pseudo_F10, Pseudo_YMC11/02/R656 and Pseudo_JD024 in BK3, Pseudo_MD8 in BK4 and Pseudo_Phi3 in BK5, suggesting that they can contribute to the keratitis virulence.

Further, to evaluate the role of a CRISPR-Cas system (Clustered Regularly Interspaced Short Palindromic Repeats and Crisper associated genes) in resistance, we investigated the genes of the CRISPR-Cas system. The CRISPR loci and CRISPR-associated proteins Csy1, Csy2, Csy3 and Csy4 were detected only in MDR strain BK6, which was cultured from patients with good clinical outcomes. The results suggest that the CRISPR-Cas system may not have a direct role in the clinical outcome.

## 4. Conclusions

In summary, we showed a large genome size and more genes in the MDR group than the non-MDR group. However, 90% of the virulence genes were detected in all strains. Among them, exoU and exoS of the T3SS, the main contributor for keratitis pathogenesis, was detected in MDR and non-MDR strains, respectively. Previous studies reported that MDR strains with exoU might lead to poor clinical outcomes [11, 40]; however, in this study, one MDR strain carrying exoU was isolated from a patient with good clinical outcomes. Despite the above, the strain-specific genes responsible for virulence and drug-resistance were detected in the poor outcome group. Of the strain-specific genes, fliC and fliD reported to altering the host immune response, suggesting their role in clinical outcome. However, further studies are required to confirm their role.

## Ethics approval and consent to participate

This study adhered to the tenets of the Declaration of Helsinki and ethics committee approval was obtained from the Institutional Review Board of the Aravind Eye Care System. Informed consent was obtained from each patient before surgery.

## Acknowledgements

This work was supported by Aravind Eye Hospital, Madurai.

## Declaration of competing interest

The authors declare that they have no known competing financial interests or personal relationships that could have appeared to influence the work reported in this paper.

## 1. Supplementary Figures

**1.1 Fig S1.**
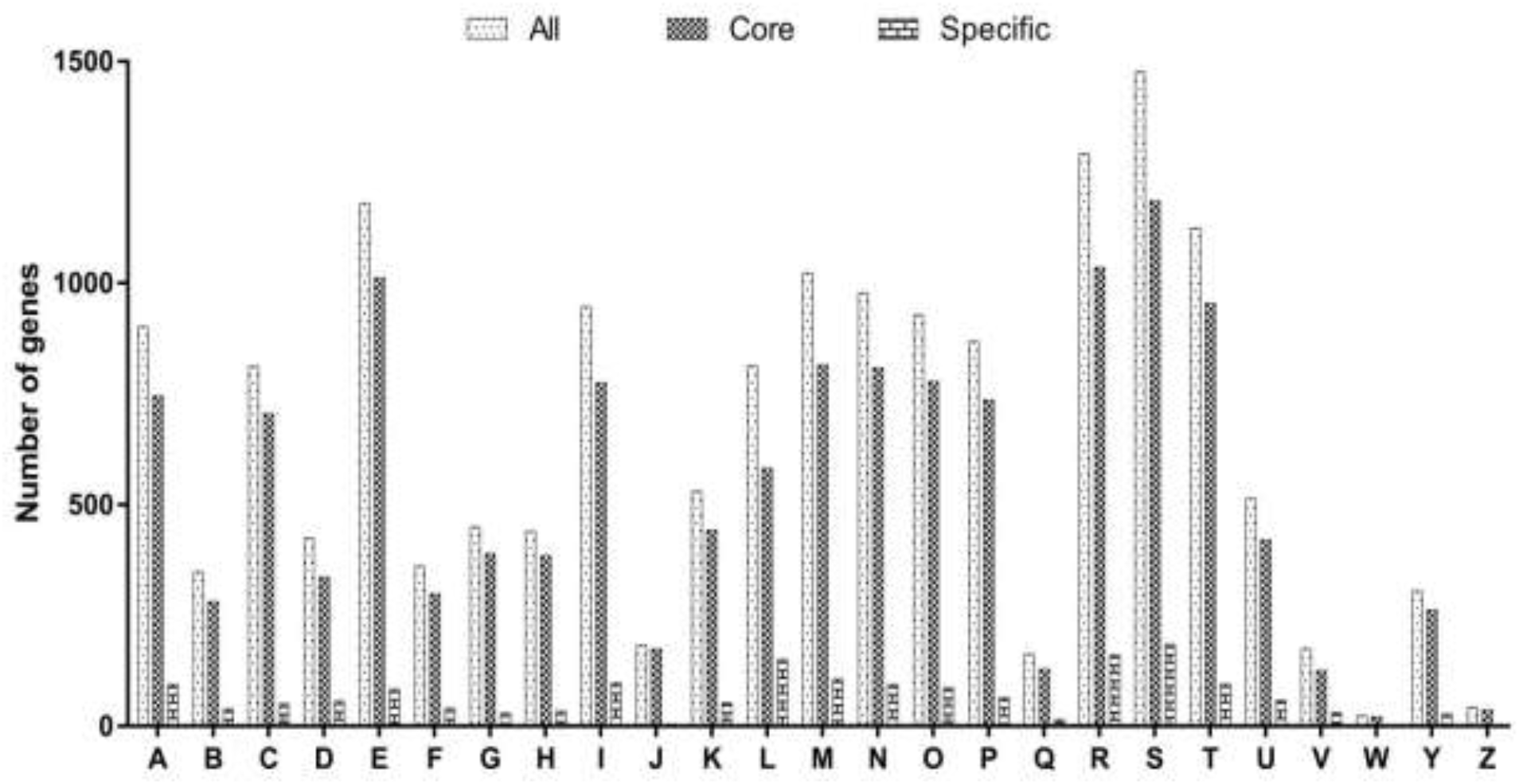
The distribution of genes according to the COG classification. Y-axis indicates the number of genes in different COG categories. (A) RNA processing and modification; (B) Chromatin structure and dynamics; (C) Energy production and conversion; (D) Cell cycle control, mitosis, and meiosis; (E) Amino acid transport and metabolism; (F) Nucleotide transport and metabolism; (G) Carbohydrate transport and metabolism; (H) Coenzyme transport and metabolism; (I) Lipid transport and metabolism; (J) Translation, ribosomal structure and biogenesis; (K) Transcription; (L) Replication, recombination, and repair; (M) Cell wall/membrane/envelope biogenesis; (N) Cell motility; (O) Post-translational modification, protein turnover, chaperones; (P) Inorganic ion transport and metabolism; (Q) Secondary metabolites biosynthesis, transport, and catabolism; (R) General function prediction only; (S) Function unknown; (T) Signal transduction mechanisms; (U) Intracellular trafficking and secretion; (V) Defense mechanisms; (W) Extracellular structures; (Y) Nuclear structure; (Z) Cytoskeleton.

## 2. Supplementary Tables

**2.1 Table S1.**
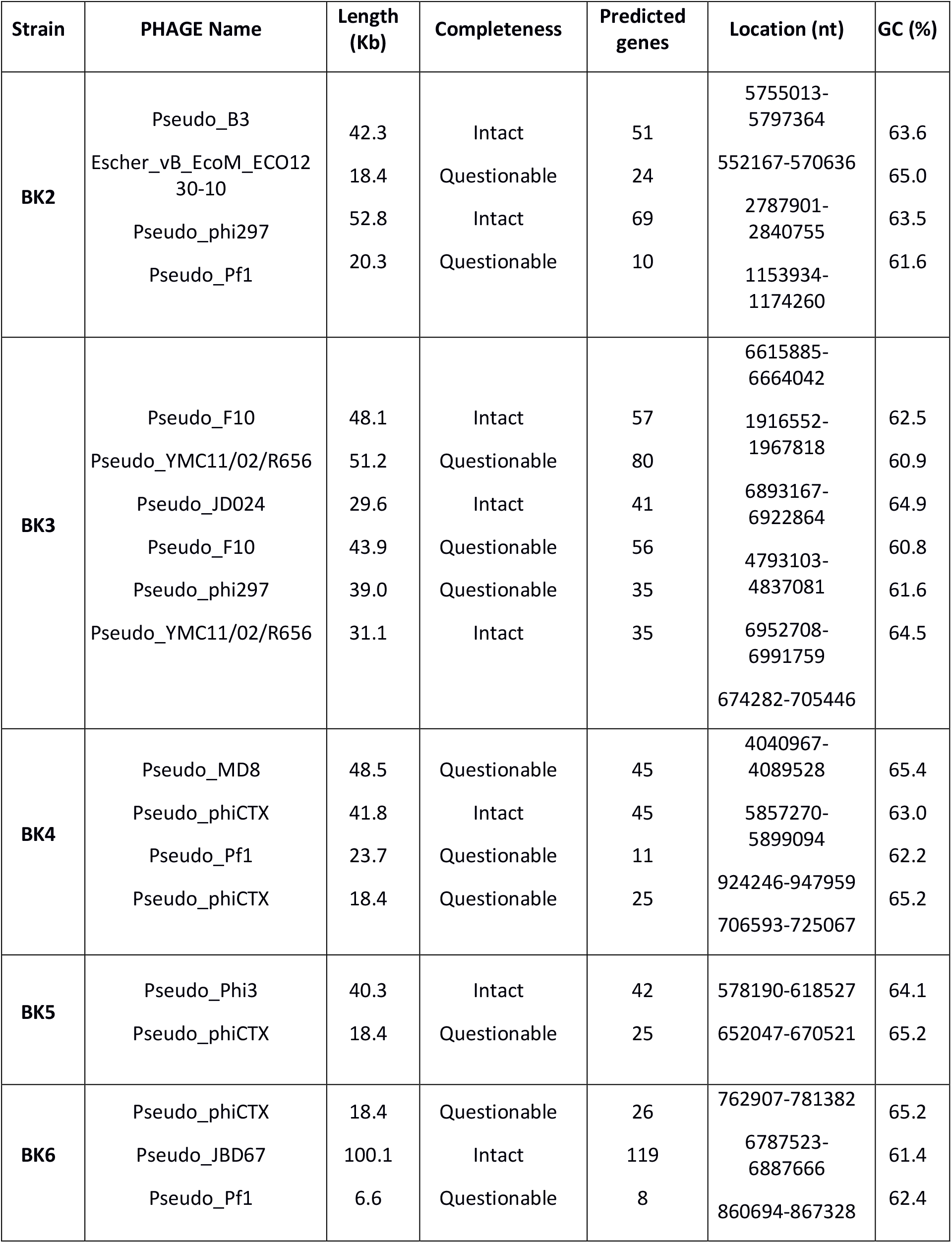
Prophages identified in ocular *P. aeruginosa* genomes.

**2.2 Table S2.**
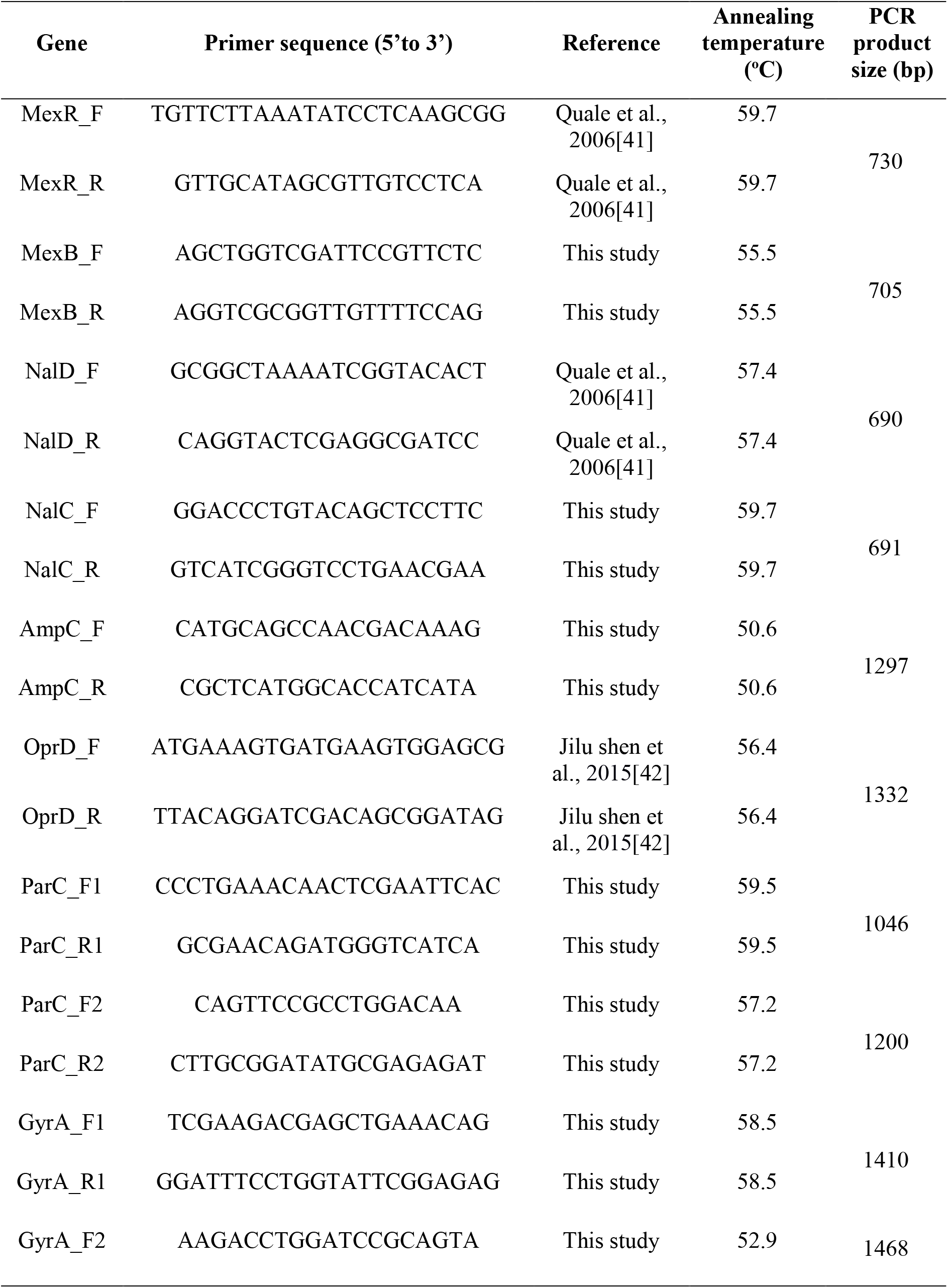
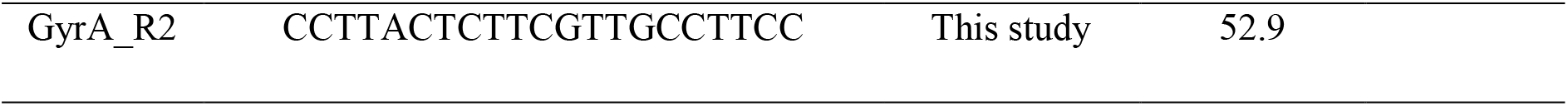
Primer sequences, annealing temperatures and amplicon size of targeted genes for Sanger sequencing.

## 3. Primers sequences for identification of Type 3 secretion system (T3SS) genotype

The primer sequences for identification of T3SS by qRT-PCR are as follows: *exoS*, FP (5’CCCGGAACTGCGGCGCGAAATCAC3’); RP (5’CCGGT TTGCTTGCCAGGTCGAGAG3’); *exoT*, FP (5’ACTCCCCGGGAGGCGCAACAAC3’) RP (5’CTTGTCGACAGGCTCGCCCTTT3’) and *exoU*, FP (5’CAGGAGCGAGTCGGTG AACATCTG3’); RP (5’CGCCCGACGTAACAGAGCTAC3’).

## Notes

### Competing Interest Statement

The authors have declared no competing interest.

## References

[1] T. Rattanatam, W.J. Heng, C.J. Rapuano, P.R. Laibson, E.J. Cohen, Trends in Contact Lens–related Corneal Ulcers, Cornea, 20 (2001) 290–294.

[2] D.E. Bradley, Basic characterization of a *Pseudomonas aeruginosa* pilus-dependent bacteriophage with a long noncontractile tail, Journal of Virology, 12 (1973) 1139–1148.

[3] V. Pelicic, Type IV pili: e pluribus unum?, Molecular Microbiology, 68 (2008) 827–837.

[4] R. O’Callaghan, A. Caballero, A. Tang, M. Bierdeman, *Pseudomonas aeruginosa* Keratitis: Protease IV and PASP as Corneal Virulence Mediators, Microorganisms, 7 (2019) 281.

[5] J. Lomholt, K. Poulsen, M. Kilian, Epidemic Population Structure of *Pseudomonas aeruginosa*: Evidence for a Clone That Is Pathogenic to the Eye and That Has a Distinct Combination of Virulence Factors, Infection and Immunity, 69 (2001) 6284–6295.

[6] E. Lee, Role of *Pseudomonas aeruginosa* ExsA in Penetration through Corneal Epithelium in a Novel In Vivo Model, Investigative Ophthalmology & Visual Science, 44 (2003) 5220–5227.

[7] A. M, L. D, Pseudomonas Keratitis, a Review of Where We’ve Been and What Lies Ahead, Journal of Microbial & Biochemical Technology, 08 (2015).

[8] V. Finck-Barbançon, J. Goranson, L. Zhu, T. Sawa, J.P. Wiener-Kronish, S.M.J. Fleiszig, C. Wu, L. Mende-Mueller, D.W. Frank, ExoU expression by *Pseudomonas aeruginosa* correlates with acute cytotoxicity and epithelial injury, Molecular Microbiology, 25 (1997) 547–557.

[9] R. Stewart, L. Wiehlmann, K. Ashelford, S. Preston, E. Frimmersdorf, B. Campbell, T. Neal, N. Hall, S. Tuft, S. Kaye, C. Winstanley, Genetic Characterization Indicates that a Specific Subpopulation of *Pseudomonas aeruginosa* Is Associated with Keratitis Infections, Journal of clinical microbiology, 49 (2011) 993–1003.

[10] M.C. Wolfgang, B.R. Kulasekara, X. Liang, D. Boyd, K. Wu, Q. Yang, C.G. Miyada, S. Lory, Conservation of genome content and virulence determinants among clinical and environmental isolates of *Pseudomonas aeruginosa*, Proceedings of the National Academy of Sciences, 100 (2003) 8484.

[11] N. Murugan, J. Malathi, V. Umashankar, H.N. Madhavan, Resistome and pathogenomics of multidrug resistant (MDR) *Pseudomonas aeruginosa* VRFPA03, VRFPA05 recovered from alkaline chemical keratitis and post-operative endophthalmitis patient, Gene, 578 (2016) 105–111.

[12] A. Darling, B. Mau, N. Perna, Darling AE, Mau B, Perna NT.. progressiveMauve: multiple genome alignment with gene gain, loss and rearrangement. PLoS ONE 5: e11147, PloS one, 5 (2010) e11147.

[13] T. Brettin, J.J. Davis, T. Disz, R.A. Edwards, S. Gerdes, G.J. Olsen, R. Olson, R. Overbeek, B. Parrello, G.D. Pusch, M. Shukla, J.A. Thomason, R. Stevens, V. Vonstein, A.R. Wattam, F. Xia, RASTtk: A modular and extensible implementation of the RAST algorithm for building custom annotation pipelines and annotating batches of genomes, Scientific Reports, 5 (2015).

[14] A.J. Page, C.A. Cummins, M. Hunt, V.K. Wong, S. Reuter, M.T.G. Holden, M. Fookes, D. Falush, J.A. Keane, J. Parkhill, Roary: rapid large-scale prokaryote pan genome analysis, Bioinformatics, 31 (2015) 3691–3693.

[15] M.V. Larsen, S. Cosentino, S. Rasmussen, C. Friis, H. Hasman, R.L. Marvig, L. Jelsbak, T. Sicheritz-Pontén, D.W. Ussery, F.M. Aarestrup, O. Lund, Multilocus Sequence Typing of Total-Genome-Sequenced Bacteria, Journal of clinical microbiology, 50 (2012) 1355.

[16] E. Zankari, H. Hasman, S. Cosentino, M. Vestergaard, S. Rasmussen, O. Lund, F. Aarestrup, M. Larsen, Identification of Acquired Antimicrobial Resistance Genes, The Journal of antimicrobial chemotherapy, 67 (2012) 2640–2644.

[17] L. Chen, J. Yang, J. Yu, Z. Yao, L. Sun, Y. Shen, Q. Jin, Chen L.. VFDB: a reference database for bacterial virulence factors. Nucleic Acids Res 33: D325-D328, Nucleic acids research, 33 (2005) D325–328.

[18] Y. Zhou, Y. Liang, K.H. Lynch, J.J. Dennis, D.S. Wishart, PHAST: A Fast Phage Search Tool, Nucleic Acids Research, 39 (2011) W347–W352.

[19] D. Couvin, A. Bernheim, C. Toffano-Nioche, M. Touchon, J. Michalik, B. Néron, E.P.C. Rocha, G. Vergnaud, D. Gautheret, C. Pourcel, CRISPRCasFinder, an update of CRISRFinder, includes a portable version, enhanced performance and integrates search for Cas proteins, Nucleic acids research, 46 (2018) W246–W251.

[20] T.J. Treangen, B.D. Ondov, S. Koren, A.M. Phillippy, The Harvest suite for rapid core-genome alignment and visualization of thousands of intraspecific microbial genomes, Genome Biology, 15 (2014) 524.

[21] D. Subedi, A. Vijay, G. Kohli, S. Rice, M. Willcox, Comparative genomics of clinical strains of *Pseudomonas aeruginosa* strains isolated from different geographic sites, Scientific Reports, 8 (2018).

[22] B. Valot, C. Guyeux, J.Y. Rolland, K. Mazouzi, X. Bertrand, D. Hocquet, What It Takes to Be a *Pseudomonas aeruginosa*? The Core Genome of the Opportunistic Pathogen Updated, PloS one, 10 (2015) e0126468.

[23] D. Girlich, T. Naas, P. Nordmann, Biochemical Characterization of the Naturally Occurring Oxacillinase OXA-50 of *Pseudomonas aeruginosa*, Antimicrobial Agents and Chemotherapy, 48 (2004) 2043–2048.

[24] Z. M. Jaafar, M. Dhahi, K. Abdul, M. Safaa, Molecular identification and antibiotics resistance genes profile of *Pseudomonas aeruginosa* isolated from Iraqi patients, African Journal of Microbiology Research, 8 (2014) 2183–2192.

[25] P.A. White, H.W. Stokes, K.L. Bunny, R.M. Hall, Characterisation of a chloramphenicol acetyltransferase determinant found in the chromosome of *Pseudomonas aeruginosa*, FEMS Microbiology Letters, 175 (1999) 27–35.

[26] K. Thirumalmuthu, B. Devarajan, L. Prajna, V. Mohankumar, Mechanisms of Fluoroquinolone and Aminoglycoside Resistance in Keratitis-Associated *Pseudomonas aeruginosa*, Microbial Drug Resistance, 25 (2019) 813–823.

[27] V. Tam, A. Schilling, M. LaRocco, L. Gentry, K. Lolans, J. Quinn, K. Garey, Prevalence of AmpC over-expression in bloodstream isolates of *Pseudomonas aeruginosa*, Clinical microbiology and infection: the official publication of the European Society of Clinical Microbiology and Infectious Diseases, 13 (2007) 413–418.

[28] G. Cabot, S. Bruchmann, X. Mulet, L. Zamorano, B. Moya, C. Juan, S. Haussler, A. Oliver, *Pseudomonas aeruginosa* Ceftolozane-Tazobactam Resistance Development Requires Multiple Mutations Leading to Overexpression and Structural Modification of AmpC, Antimicrobial Agents and Chemotherapy, 58 (2014).

[29] J. Lee, Y. Lee, Y. Park, B.-S. Kim, Alterations in the GyrA and GyrB Subunits of topoisomerase II and the ParC and ParE subunits of topoisomerase IV in ciprofloxacin-resistant clinical isolates of *Pseudomonas aeruginosa*, International journal of antimicrobial agents, 25 (2005) 290–295.

[30] R. Salma, F. Dabboussi, I. al Kassaa, R. Khudary, M. Hamze, GyrA and parC mutations in quinolone-resistant clinical isolates of *Pseudomonas aeruginosa* from Nini Hospital in north Lebanon, Journal of infection and chemotherapy: official journal of the Japan Society of Chemotherapy, 19 (2012).

[31] H. Mouneimné, J. Robert, V. Jarlier, E. Cambau, Type II topoisomerase mutations in ciprofloxacin-resistant strains of *Pseudomonas aeruginosa*, Antimicrobial Agents and Chemotherapy, 43 (1999) 62–66.

[32] M.M. Ochs, M.P. McCusker, M. Bains, R.E. Hancock, Negative regulation of the *Pseudomonas aeruginosa* outer membrane porin OprD selective for imipenem and basic amino acids, Antimicrobial Agents and Chemotherapy, 43 (1999) 1085–1090.

[33] M.L. Sobel, D. Hocquet, L. Cao, P. Plesiat, K. Poole, Mutations in PA3574 (nalD) lead to increased MexAB-OprM expression and multidrug resistance in laboratory and clinical isolates of *Pseudomonas aeruginosa*, Antimicrobial Agents and Chemotherapy, 49 (2005) 1782–1786.

[34] D. Ince, F. Sutterwala, T. Yahr, Secretion of Flagellar Proteins by the *Pseudomonas aeruginosa* Type III Secretion-Injectisome System, Journal of bacteriology, 197 (2015).

[35] N. Dasgupta, M. Wolfgang, A. Goodman, S. Arora, J. Jyot, S. Lory, R. Ramphal, Four-tiered transcriptional regulatory circuit controls flagellar biogenesis in *Pseudomonas aeruginosa*, Molecular Microbiology, 50 (2003) 809–824.

[36] S.L.N. Kilmury, L.L. Burrows, The *Pseudomonas aeruginosa* PilSR Two-Component System Regulates Both Twitching and Swimming Motilities, mBio, 9 (2018) e01310–01318.

[37] D. Nunn, S. Bergman, S. Lory, Products of Three Accessory Genes, pilB, pilC, and pilD, are Required for Biogenesis of *Pseudomonas aeruginosa* Pili, Journal of bacteriology, 172 (1990) 2911–2919.

[38] H.K. Takhar, K. Kemp, M. Kim, P.L. Howell, L.L. Burrows, The platform protein is essential for type IV pilus biogenesis, J Biol Chem, 288 (2013) 9721–9728.

[39] P. Knezevic, M. Voet, R. Lavigne, Prevalence of Pf1-like (pro)phage genetic elements among *Pseudomonas aeruginosa* isolates, Virology, 483 (2015) 64–71.

[40] D. Borkar, N. Acharya, C. Leong, P. Lalitha, M. Srinivasan, C. Oldenburg, V. Cevallos, T. Lietman, D. Evans, S. Fleiszig, Cytotoxic clinical isolates of *Pseudomonas aeruginosa* identified during the Steroids for Corneal Ulcers Trial show elevated resistance to fluoroquinolones, BMC ophthalmology, 14 (2014) 54.

## References for SI

[41] J. Quale, S. Bratu, J. Gupta, D. Landman, Interplay of efflux system, ampC, and oprD expression in carbapenem resistance of Pseudomonas aeruginosa clinical isolates, Antimicrobial Agents and Chemotherapy 50 (2006) 1633–1641.

[42] J. Shen, Y. Pan, Y. Fang, Role of the Outer Membrane Protein OprD2 in Carbapenem-Resistance Mechanisms of Pseudomonas aeruginosa, PloS one 10 (2015) e0139995.

